# Diversity, taxonomy and evolution of archaeal viruses of the class *Caudoviricetes*

**DOI:** 10.1101/2021.05.26.445827

**Authors:** Ying Liu, Tatiana A. Demina, Simon Roux, Pakorn Aiewsakun, Darius Kazlauskas, Peter Simmonds, David Prangishvili, Hanna M. Oksanen, Mart Krupovic

## Abstract

The archaeal tailed viruses (arTV), evolutionarily related to tailed double-stranded DNA bacteriophages of the class *Caudoviricetes*, represent the most common isolates infecting halophilic archaea. Only a handful of these viruses have been genomically characterized, limiting our appreciation of their ecological impacts and evolution. Here, we present 37 new genomes of haloarchaeal tailed virus isolates, more than doubling the current number of sequenced arTVs. Analysis of all 63 available complete genomes of arTVs, which we propose to classify into 14 new families, suggests ancient divergence of archaeal and bacterial tailed viruses and points to an extensive sharing of genes involved in DNA metabolism and counter defense mechanisms, illuminating common strategies of virus-host interactions with tailed bacteriophages. Coupling of the comparative genomics with the host range analysis on a broad panel of haloarchaeal species uncovered four distinct groups of viral tail fiber adhesins controlling the host range expansion. The survey of metagenomes using viral hallmark genes suggests that the global architecture of the arTV community is shaped through recurrent transfers between different biomes, including hypersaline, marine and anoxic environments.

## INTRODUCTION

Bacteriophages with helical tails and icosahedral capsids (tailed bacteriophages), classified into the class *Caudoviricetes* ^1^, represent the most widespread, abundant and diverse group of viruses on our planet ^2^, and are likely to infect hosts from most, if not all, known bacterial lineages ^3^. Extensive experimental studies conducted over several decades, coupled with comparative analysis of several thousands of complete tailed phage genomes currently available in the public sequence databases, have resulted in detailed understanding of the mechanisms that govern the biology, ecology and evolution of this group of viruses ^2,4-6^. Due to their ubiquity, tailed bacteriophages have a profound impact on the functioning of the biosphere through regulating the structure, composition and dynamics of bacterial populations in diverse environments, from marine ecosystems to the human gut, and modulate major biogeochemical cycles ^7-9^. Archaeal tailed viruses (arTV) are morphologically indistinguishable from tailed bacteriophages ^1,4,10-12^. Similar to their bacterial relatives, the helical tails of arTVs can be short (podovirus morphology), long non-contractile (siphovirus morphology) or contractile (myovirus morphology) ^13-15^.

The arTVs have been thus far isolated on halophilic (class *Halobacteria*) and methanogenic (family *Methanobacteriaceae*) archaea, both belonging to the phylum Euryarchaeota ^16-25^. Related proviruses have also been sighted in other lineages of the Euryarchaeota as well as in ammonia-oxidizing Thaumarchaeota and Aigarchaeota ^26-30^, whereas recent metagenomics studies revealed novel groups of arTVs putatively infecting marine group II Euryarchaeota, Thaumarchaeota and Thermoplasmata ^31-35^. Ecologically, it has been shown that virus-mediated lysis of archaea in the deep ocean is more rapid than that of bacteria, suggesting an important ecological role of archaeal viruses in marine ecosystems ^36^. Evolutionarily, the broad distribution of tailed (pro)viruses in both bacteria and archaea suggests that viruses of this type were part of the virome associated with the last universal cellular ancestor, LUCA 3.

Genomic and structural analyses have shown that archaeal and bacterial tailed viruses have similar genomic organizations, with genes clustered into functional modules, and share homologous virion morphogenesis modules, including the major capsid protein (MCP) with the characteristic HK97 fold and the genome packaging terminase complex, suggesting common principles of virion assembly ^11,13,22,30,37,38^. Nevertheless, at the sequence level, arTVs are strikingly diverse showing little similarity to each other and virtually no recognizable similarity to their bacterial relatives, indicative of scarce sampling of the archaeal virosphere ^38^. Indeed, for several thousands of complete genome sequences of tailed bacteriophages ^2^, only 25 arTV isolates have been sequenced thus far. The low number of isolates severely limits our appreciation of the ecological impacts of arTVs and obscures the evolutionary history of this important and ancient group of viruses.

Virus discovery in the “age of metagenomics” is increasingly performed by culture-independent methods, whereby viral genomes are sequenced directly from the environment. This is a powerful approach which has already yielded thousands of viral genomes, providing unprecedented insights into virus diversity, environmental distribution and evolution ^39-46^. The limitation of viral metagenomics, however, is that the exact host species for the sequenced viruses typically remain unknown and many molecular aspects of virus-host interactions cannot be accurately predicted. Here, to further explore the biology and diversity of arTVs, we sequenced the genomes of 37 viruses which infect different species of halophilic archaea and were isolated from hypersaline environments using classical approaches ^16,17^. Collectively, our results provide the first global overview of arTV diversity and evolution and establish a robust taxonomic framework for their classification.

## RESULTS AND DISCUSSION

### Overview of new haloarchaeal tailed viruses

We sequenced a collection of 37 arTVs (5 siphoviruses and 32 myoviruses) infecting haloarchaeal species belonging to the genera *Halorubrum* and *Haloarcula* ^16,17^, more than doubling the number of complete genomes of arTVs (Table S1). The sequenced viruses originate from geographically remote locations, including Thailand, Israel, Italy and Slovenia, and in combination with the previously described isolates, provide a substantially improved genomic insight into the global distribution of arTVs. The viruses possess double-stranded DNA (dsDNA) genomes ranging from 35.3 to 104.7 kb in length. Genomes of several isolates were nearly identical (<17 nucleotide polymorphisms; Table S1) and analysis of these genomes, in combination with host range experiments, was particularly illuminating towards the host range evolution among halophilic arTVs (see below).

### Archaeal tailed viruses represent a distinct group within the prokaryotic virosphere

To assess the global diversity of arTVs and analyze their relationship to bacterial members of the class *Caudoviricetes*, we supplemented the 37 genomes sequenced herein with the genomes of prokaryotic viruses available in GenBank and analyzed the dataset using GRAViTy ^47,48^ and vConTACT v2.0 ^49^. The combined dataset included 63 complete arTV genomes (Table S1), three of which were from viruses infecting methanogenic archaea, whereas all others were from haloarchaeal viruses. The GRAViTy tool classifies viruses into family-level taxonomic groupings according to homology between viral genes and similarities in genome organizations, which are expressed using composite generalized Jaccard (CGJ) distances ^47,48^. We used a CGJ distance of over 0.8 as the threshold for family-level assignment, consistent with the family-level classification for eukaryotic viruses and the recently created families of bacterial viruses ^47,48,50^. GRAViTy analysis of the global prokaryotic virome classified arTVs into two large assemblages, which could be further subdivided into 14 family-level groupings (CGJ distance ≥ 0.8) (Fig. 1). To reveal a finer taxonomic structure within the archaeal tailed virus assemblage, we relied on the network analytics implemented in vConTACT v2.0, which has been specifically developed and calibrated to identify genus-level groupings of prokaryotic viruses ^49^. Consistent with the GRAViTy results, the network analysis revealed two assemblages of arTVs, which were disconnected from all known bacterial and non-tailed archaeal viruses (Fig. 2). Viruses within the two clades formed 23 genus-level and 14 family-level groups, many containing just one or two members, indicating that genetic diversity of archaeal viruses remains largely undersampled. The family-level and genus-level groupings are described in Supplementary text.

**Fig. 1.**
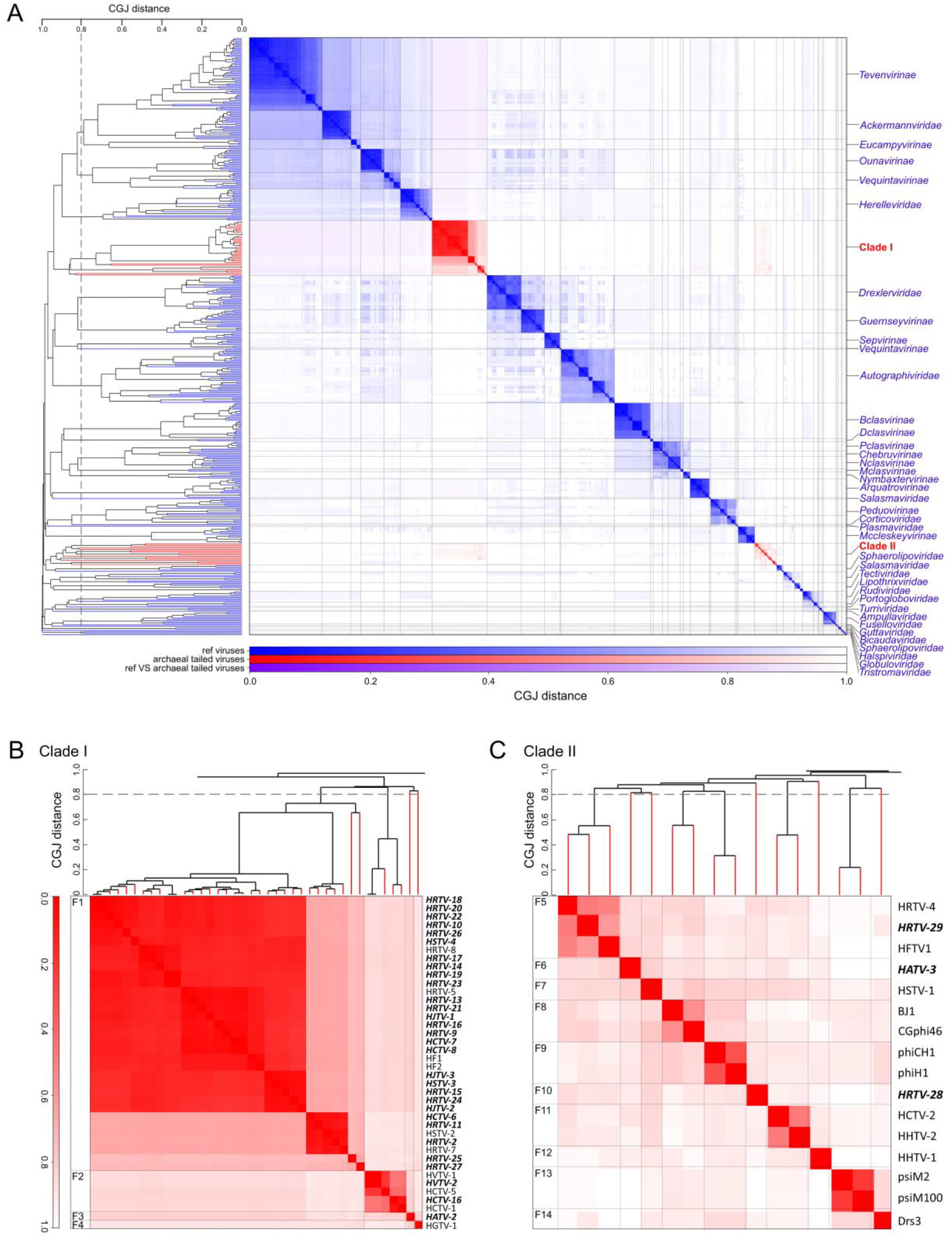
Genome relationships between prokaryotic dsDNA viruses. (A) Heat map and dendrogram of composite generalized Jaccard (CGJ) distances for classified bacteriophages and archaeal dsDNA viruses. Branches and clusters corresponding to arTVs are shown in red, whereas those of other viruses are in blue. (B) Zoom in on the arTV Clade I. (C) Zoom in on the arTV Clade II. Viruses sequenced in this study are highlighted in bold. CGJ distance of 0.8, chosen as a family-level threshold, is indicated with a broken line, with the family-level groups (F1-F14) indicated on the left of the heatmap.

**Fig. 2.**
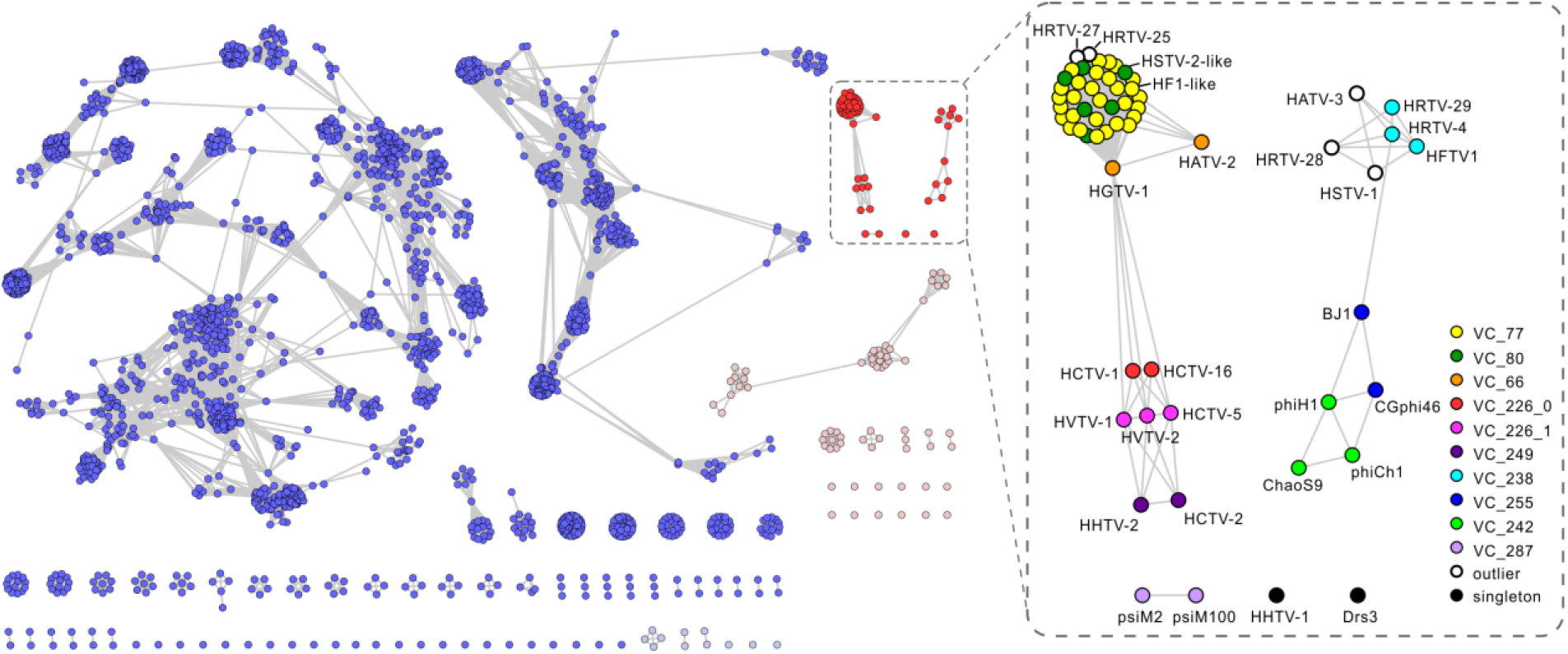
The network-based analysis of shared protein clusters (PCs) among arTVs and the prokaryotic dsDNA viruses. The nodes represent viral genomes, and the edges represent the strength of connectivity between each genome based on shared PCs. Nodes representing genomes of arTVs are in red and other archaeal viruses are in light red, whereas those representing genomes of tailed bacteriophages are in blue and other bacteriophages are in light blue (left panel). The viral clusters (VCs) of arTVs are enlarged and labeled in the right panel.

Clade I in the GRAViTy analysis forms a sister branch to several groups of bacterial myoviruses, including families *Ackermannviridae, Herelleviridae* and T4-like bacteriophages, and consists of four family-level groups (Fig. 1A, B; Supplementary text), with family (F) 1 being by far the largest, with 39 members, which can be further divided into four genus-level (G) subgroups (Fig. 2). Clade I is a cohesive assemblage consisting exclusively of haloarchaeal viruses and held together by 33 protein clusters, including those involved in virion morphogenesis, genome replication and repair, nucleotide metabolism, and several proteins of unknown function (Table S3). By contrast, Clade II is less cohesive and consists of 10 family-level groupings, half of which consist of singletons (Fig. 1A, C), and includes viruses of both halophilic and methanogenic archaea. Viruses from different family-level groups in this clade share an overlapping set of genes (52 protein clusters), including those involved in virion morphogenesis, genome replication and repair, DNA metabolism and several other functionally diverse proteins. However, unlike in Cluster I, the set of genes holding the Cluster II together, displays sporadic distribution, typically connecting 2-3 virus families (Table S3).

Based on the results of comprehensive comparative genomics analysis (see Supplementary text), we propose classifying all known arTVs into 14 new families. The families *Hafunaviridae, Soleiviridae, Halomagnusviridae*, and *Pyrstoviridae* include viruses with icosahedral heads and long contractile tails (myovirus morphotype), whereas the families *Queuoviridae, Haloferuviridae, Flexireviridae, Vertoviridae, Suolaviridae, Saparoviridae, Madisaviridae, Leisingerviridae*, and *Anaerodiviridae* contain viruses with icosahedral heads and long non-contractile tails (siphovirus morphotype). The *Shortaselviridae* is the only family of viruses with short tails (podovirus morphology). The names of the 23 proposed genera are listed in Table S1. Members of the same proposed genus typically share more than 60% of their proteins and members of the same family share 20-50% of homologous proteins, whereas viruses from different families share less than 10% of proteins (Fig. 3, Fig. S2).

**Fig. 3.**
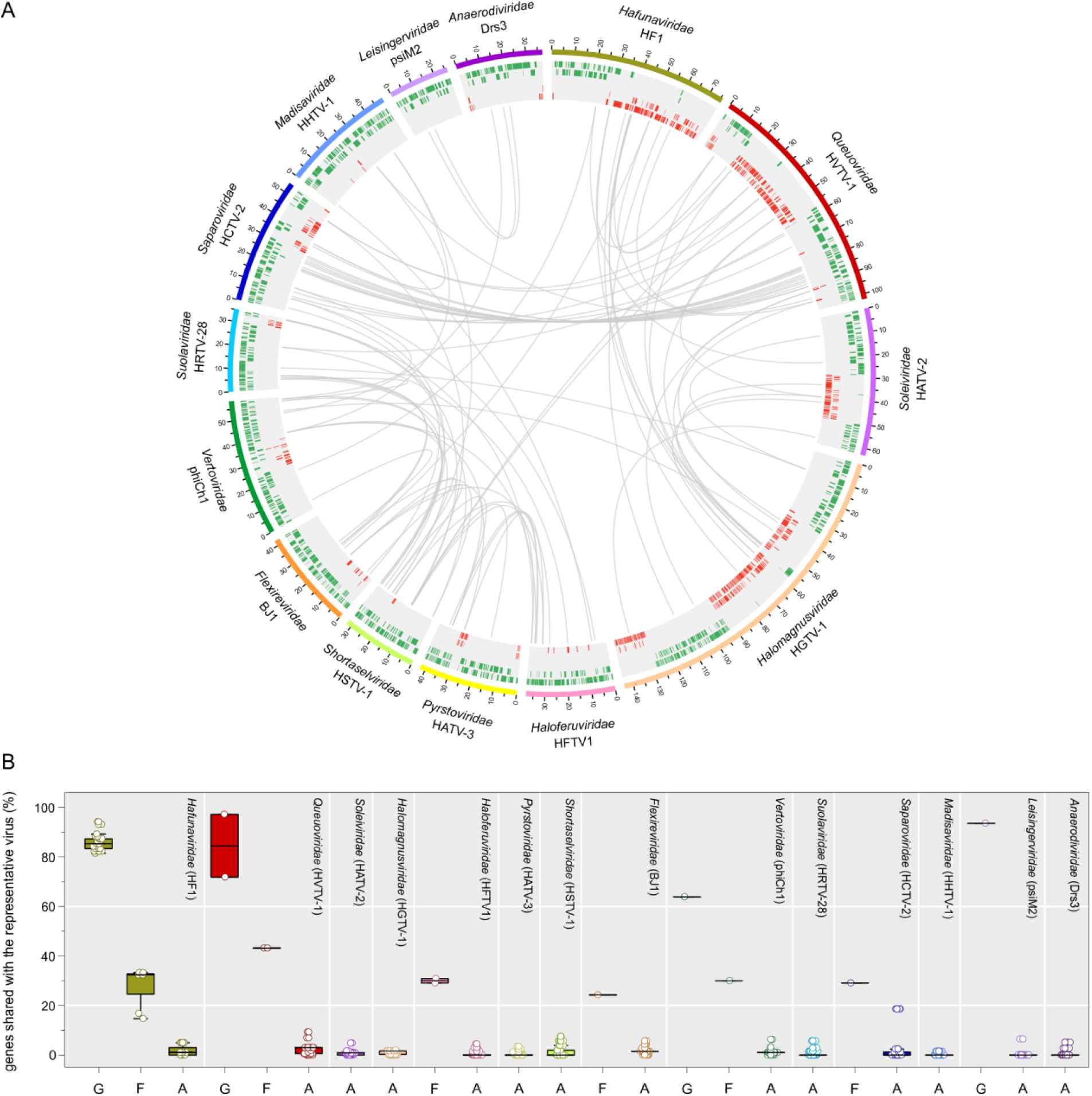
Overview of homologous proteins shared by arTVs. (A) The circus plot displays gene content similarity between representative members of each proposed family. Viral genomes from different families are indicated with distinct colors. The genomic coordinates (kb) are indicated on genome maps. Putative ORFs distributed on five tracks (to avoid overlapping) are represented by green and red tiles on the forward and reverse strands, respectively. Homologous proteins are linked by grey lines. The representative viruses and the proposed virus family names are indicated. (B) The box plot shows the percentage of genes shared by a representative virus from each family with other members of the same genus (G) and family (F) as well as with arTVs from other families (A). Each box represents the middle 50th percentile of the data set and is derived using the lower and upper quartile values. The median value is displayed by a horizontal line. Whiskers represent the maximum and minimum values with the range of 1.5 IQR. Each virus is represented by a dot. In the circus and box plots analyses, proteins with over 30% amino acid sequence identity and E-value < 1 × 10^−25^ in our arTV database are considered to be homologous.

### Gene content of archaeal tailed viruses

All virus genes were functionally annotated using sensitive profile-profile hidden Markov model (HMM) comparisons using HHpred ^51^ (Table S5). The virus-encoded proteins were further classified into functional categories based on their affiliation to the archaeal clusters of orthologous genes (arCOG) ^52^ (Table S5). Apart from the “Virus related” proteins, the most frequent functional category assigned to archaeal virus proteins was the “Information storage and processing”, with the “Genome replication, recombination and repair” subcategory being most strongly enriched (Fig. 4A). Proteins of the “Defense mechanisms” subcategory from the “Cellular processes and signaling” category, primarily including diverse nucleases and DNA methyltransferases (MTases), were also abundant. Finally, a substantial fraction of proteins was assigned to the “Metabolism” category, with proteins involved in nucleotide transport and metabolism being most common (Fig. 4A). Below we highlight some of the observations with the more detailed description provided in the Supplementary text.

**Fig 4.**
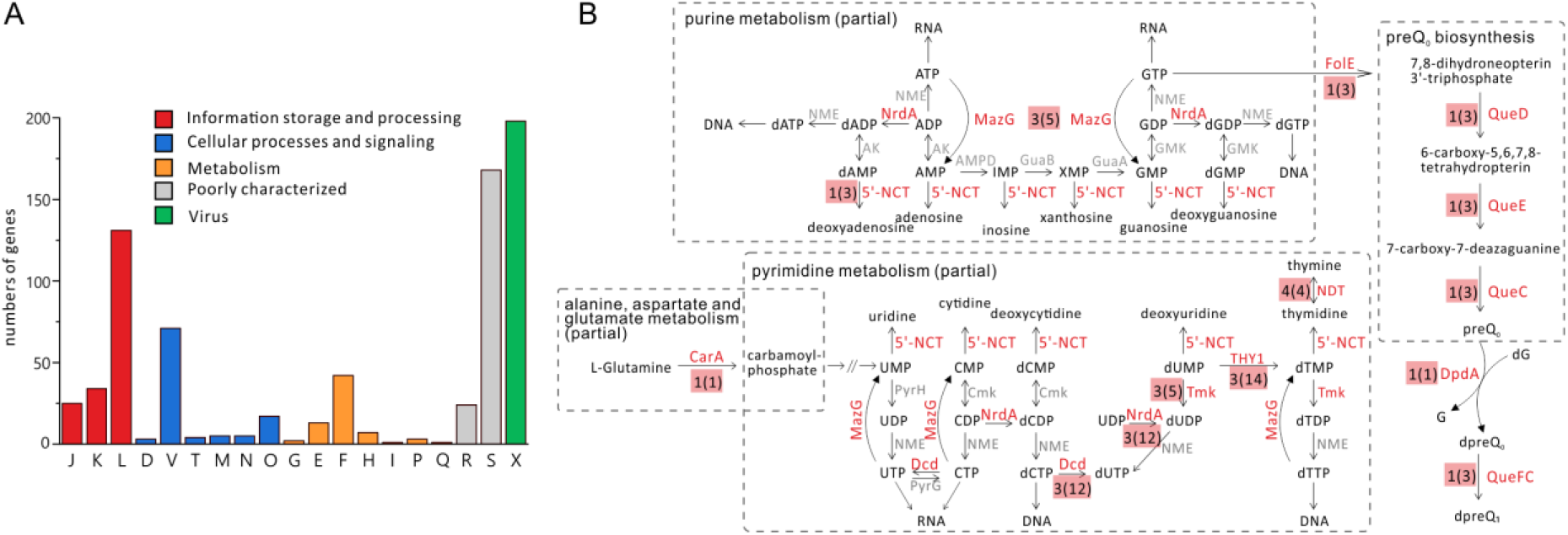
Classification of genes encoded by arTVs. (A) Classification of arTV genes into arCOG functional categories. The homologous gene shared by viruses in the same species is counted as one. The letters represent: J, translation, ribosomal structure and biogenesis; K, transcription; L, replication, recombination and repair; D, cell cycle control, cell division, chromosome partitioning; V, defense mechanisms; T, signal transduction mechanisms; M, cell wall/membrane/envelope biogenesis; N, cell motility; O, posttranslational modification, protein turnover, chaperons; G, carbohydrate transport and metabolism; E, amino acid transport and metabolism; F, nucleotide transport and metabolism; H, coenzyme transport and metabolism; I, lipid transport and metabolism; P, inorganic ion transport and metabolism; Q, secondary metabolites biosynthesis, transport and catabolism; R, general function prediction only; S, function unknown; X, virus-related. (B) Schematic showing nucleotide-related (partial) metabolic pathways, with a particular enzyme indicated for each reaction. Enzymes that are found encoded by arTVs are in red, otherwise in grey. The number of virus families that harbor the enzyme was labeled beside the name the enzyme, with the number of virus species shown in the brackets. NrdA: ribonucleotide diphosphate reductase, MazG: NTP pyrophosphatase, 5’-NCT: 5’-deoxyribonucleotidase, CarA: Carbamoylphosphate synthase small subunit, Dcd: dCTP deaminase, Tmk: thymidylate kinase, THY1: thymidylate synthase thyX, NDT: nucleoside deoxyribosyltransferase, FolE: GTP cyclohydrolase I, QueC: queuosine biosynthesis protein, QueD: 6-pyruvoyl tetrahydropterin synthase, QueE: 7-carboxy-7-deazaguanine synthase, DpdA: paralog of queuine tRNA-ribosyltransferase, QueFC: NADPH-dependent 7-cyano-7deazaguanine reductase.

#### Defense and counterdefense factors

Similar to tailed bacteriophages, arTVs are under strong pressure to overcome the host defenses, among which CRISPR-Cas, restriction-modification (RM) and toxin-antitoxin (TA) are the ubiquitous defense systems in both archaea and bacteria. No clear homologs of viral anti-CRISPR proteins were detected in the arTVs genomes, although several members of the *Hafunaviridae* and *Queuoviridae* encode a Cas4-like nuclease that could potentially function in counterdefense against CRISPR-Cas systems (see Supplementary text). In contrast, haloarchaeal tailed viruses (n=48) from 10 families encode diverse MTases with specificities for N6-adenine, N4-cytosine and C5-cytosine (Table S6, Fig. S3). The presence of the *MTase* genes can be linked to the presence of sequence motifs in the genomes of the corresponding viruses. For instance, GATC motif methylated by the phiCh1-like Dam MTase (G^m6^ATC) ^53^ is present at high frequency (6.47-11.04/kb) in the genomes of viruses that encode this MTase. By contrast, viruses which do not encode the Dam MTase also lack the corresponding motif in their genomes. Thus, arTVs have likely evolved to escape the host RM systems by either self-methylation via different virus-encoded MTases or purging the recognition motifs from their genomes. In addition to the stand-alone MTases, some arTVs encode accompanying restriction endonucleases (REases), forming apparently functional RM systems, which could be deployed by the viruses for degradation of the host chromosomes or genomes of competing mobile genetic elements (see Supplementary text). A similar function could be envisioned for the micrococcal nucleases hydrolyzing single- or double-stranded DNA and RNA and encoded by HRTV-17, phiH1, HCTV-2 and HHTV-2 (Table S5).

The 7-deazaguanine modifications, produced by the virus-encoded preQ0/G+ pathway, were recently detected in the genomes of diverse viruses, including HVTV-1, and shown to confer viral DNA with resistance to various type II REases ^54^. The preQ0/G+ pathway is encoded by all viruses of the *Queuoviridae* (Fig. 4B, Table S5), suggesting that these viruses depend on a similar genome modification for evasion of the host RM systems. Notably, HCTV-1 and HCTV-16 in addition encode a transporter of the queuosine precursor YhhQ. Interestingly, HRTV-29 (family *Haloferuviridae*) does not carry genes for the preQ0/G+ pathway, but encodes a DpdA homolog, a key enzyme mediating replacement of the unmodified guanine base in the DNA (Fig. 4B, Table S5). DNA modification with solely virus-encoded DpdA was demonstrated for the *Salmonella* phage 7-11 ^54^, suggesting that HRTV-29 also hijacks the host preQ0 pathway for DNA modification through its DpdA protein.

The TA systems are widely distributed in prokaryotes and have been shown to function in bacterial antiviral defense by initiating the programmed cell death, thereby preventing the virus spread ^55^. Similar to certain marine bacteriophages ^56^, viruses of the families *Queuoviridae, Vertoviridae* and *Madisaviridae* encode homologs of the nucleoside pyrophosphohydrolase MazG (Table S5), which prevents the programmed cell death by degrading the intracellular ppGpp ^57^. In addition, several arTVs encode homologs of the VapB antitoxins (arCOG08550), but not the associated toxins, suggesting a function in blocking the VapBC TA systems. Thus, arTVs appear to encode different factors for counteracting TA systems.

#### Viral metabolic genes

The pangenome of arTVs includes many predicted metabolic genes with specific or general functions (Table S7), such as phosphoadenosine phosphosulphate reductase (HRTV-29 and HFTV1) which could be involved in sulfur assimilation pathway; Class II glutamine amidotransferase (phiCh1 and ChaoS9); MaoC-like dehydratase domain protein (HATV-2) which exhibits (R)-specific enoyl-CoA hydratase activity ^58^; dual specificity protein phosphatase (HGTV-1 and viruses from the genus *Mincapvirus* of the *Hafunaviridae*), thioredoxin (HGTV-1), peroxide stress response protein YaaA (HRTV-29 and ChaoS9), ADP-ribosyltransferase (HVTV-1 and HCTV-5), sialidase (HCTV-5) and Hsp90 chaperone protein (HSTV-2 and HGTV-1). However, the most common and widespread metabolic genes in arTVs appear to be linked to the replication, transcription and translation of the viral genomes.

#### Nucleotide biosynthesis

Most haloarchaeal virus genes involved in DNA metabolism belong to the pyrimidine biosynthesis pathway and encode thymidylate synthase, thymidylate kinase, dCTP deaminase, and nucleoside deoxyribosyltransferase (Fig. 4B). These proteins are broadly encoded by viruses from families *Hafunaviridae, Queuoviridae, Soleiviridae, Halomagnusviridae, Haloferuviridae* and *Vertoviridae* (Table S7). Interestingly, HGTV-1 (family *Halomagnusviridae*) encodes the small subunit, CarA, of carbamoylphosphate (CP) synthetase, which hydrolyzes glutamine to CP, a precursor common for the biosynthesis of pyrimidines and arginine ^59^ (Fig. 4B). Thus, HGTV-1 might be redirecting the host metabolism towards de novo synthesis of pyrimidines. To our knowledge, no other viral isolate thus far has been reported to encode CarA/CarB subunits, although other enzymes in the de novo pyrimidine synthesis pathway were found encoded by bacterial viruses ^60^. Overall, the different genes identified suggest that haloarchaeal tailed viruses likely boost up the pyrimidine metabolism during infection. A similar behavior has been observed in *Pseudomonas* virus infections ^61^, suggesting that enhanced pyrimidine metabolism is critical for efficient replication of both archaeal and bacterial tailed viruses.

Viruses of the *Hafunaviridae, Queuoviridae* and *Halomagnusviridae* encode class II ribonucleotide diphosphate reductases (RnR), which convert rNTPs into corresponding dNTPs, the essential building blocks for the synthesis of viral DNA (Fig. 4B, Table S7). Viruses from the *Queuoviridae* encode putative 5’(3’)-deoxyribonucleotidases, which catalyze the dephosphorylation of nucleoside monophosphates, likely to regulate and maintain the homeostasis of nucleotide and nucleoside pools in the host cells 62 (Fig. 4B, Table S7).

Cobalamin (vitamin B12) is an important cofactor in various metabolic pathways, including DNA biosynthesis (e.g., for class II RnR) ^63^. Viruses of the *Queuoviridae* and *Saparoviridae* encode putative cobaltochelatase subunits CobS and CobT, which show sequence similarity to the two subunits encoded by certain tailed bacteriophages and cellular organisms (Table S7). CobS is an AAA+ ATPase, whereas CobT contains a von Willebrand factor type A (vWA) domain and a metal ion-dependent adhesion site (MIDAS) domain located in the C-terminal region ^64^. In HHTV-2, *cobT* is split into two ORFs with the vWA domain encoded by a separate ORF. In prokaryotes, CobS interacts with CobT and, together with the third subunit, CobN, forms an active cobaltochelatase complex ^65^. The latter catalyzes the insertion of Co^2+^ ion into the corrin ring of hydrogennobyrinate a,c-diamide, close to the final step in the aerobic cobalamin biosynthesis pathway ^63^. We found that the *cobST* two-gene cluster is widely encoded in tailed viruses that infect members of eight bacterial or archaeal orders, including Halobacteriales, as described in this study (Table S8). In cyanophages, *cobST* gene cluster is part of the core genome ^66^, although *cobT* is usually mistakenly annotated as a peptidase. Recent bioinformatic study on the genotypes of *cob* subunits in prokaryotic genomes revealed that *cobS* and *cobT* are absent in most prokaryotic genomes, including many haloarchaea, whereas *cobN* is present 65. Our finding of the broad distribution of the *cobST* gene cluster in tailed viruses, both bacterial and archaeal, suggests that viral CobST likely hijacks the host CobN, especially in the absence of the host encoded CobST, to increase the production of cobalamin, thereby promoting the virus replication.

#### Transcription and translation

It has been previously shown that arTVs encode many proteins involved in RNA metabolism ^38^ and our current study further expands the complement of viral genes implicated in RNA metabolism and protein translation. For instance, HJTV-2 (*Hafunaviridae*) encodes a Trm112 family protein, which in *Haloferax volcanii* interacts and activates two MTases, Trm9 and Mtq2, known to methylate tRNAs and release factors, respectively ^67,68^. Remarkably, four viruses, HRTV-24, HSTV-3, HJTV-3 and HRTV-15 (*Hafunaviridae*) encode homologs of the ribosomal protein L21e (Fig. S5), which mediates the attachment of the 5S rRNA onto the large ribosomal subunit and stabilizes the orientation of adjacent RNA domains ^69^. Ribosomal proteins have been recently shown to be encoded by diverse tailed bacteriophages and at least some of these proteins can be incorporated into host ribosomes ^70^. The L21e homologs identified herein are the first examples of ribosomal proteins encoded by archaeal viruses.

*Halomagnus* virus HGTV-1 encodes a record number of tRNA genes among archaeal viruses: 36 tRNA genes corresponding to all 20 proteinogenic amino acids ^38^. In addition, this virus has a number of tRNA metabolism genes: two Rnl2-family RNA ligases, which may be involved in the removal of tRNA introns ^38^; a tRNA nucleotidyl transferase (CCA-adding enzyme) involved in tRNA maturation; a Class I lysyl-tRNA synthetase which catalyzes the attachment of lysine to its cognate tRNA; and a tRNA splicing ligase RtcB possibly responsible for the repair of viral tRNAs. RtcB and several tRNAs are also encoded by viruses from the *Hafunaviridae* and *Queuoviridae* (Table S1).

### Mutations in tail fiber genes determine the broad host range of hafunaviruses

The arTVs from different families display highly variable host ranges, with some being specific to a single isolate and others infecting multiple species from different haloarchaeal genera ^16,17,22,25^. Unlike in the case of tailed bacteriophages, the factors underlying the host range in haloarchaeal viruses remain obscure. To gain insights into this question, we focused on viruses of the family *Hafunaviridae* which, with 39 myovirus isolates, is currently the largest family of arTVs. Furthermore, most hafunaviruses have broad host ranges, collectively being able to infect a dozen of different strains belonging to five haloarchaeal genera ^16,17,22^. We assessed the efficiency of plating (EOP) of 24 hafunaviruses (19 isolates from five species in the genus *Haloferacalesvirus* and five isolates from a single species in the genus *Mincapvirus*) on a panel of 24 haloarchaeal strains from five genera ^16,17^. The host ranges of viruses within each species varied considerably (Table S9, Table S10), consistent with the previous results ^16,17^.

Availability of complete genome sequences for multiple isolates from the same species allowed us to pinpoint the genes responsible for the observed differences in the host range. The most informative was the comparison of two isolates, HRTV-19 and HRTV-23, which differ by two nucleotide substitutions, but their EOPs on two of the tested strains differ by three orders of magnitude (Fig. 5, Table S9). The two mutations mapped to two adjacent genes located at the end of the tail morphogenesis module, suggesting that this locus plays a key role in determining the host specificity. This possibility is further supported by the observation that five isolates (HCTV-7, HCTV-8, HCTV-9, HCTV-10 and HCTV-11) in which the two genes are identical but sequence divergence in other loci ranges from 1 bp to ∼1.2 kb share the same host ranges, with similar EOPs (Fig. 5, Table S9). The first of the two genes encodes a glycine-rich tail fiber protein and the second gene encodes a small putative protein (Fig. S6). The glycine-rich protein displays features typical of adhesin proteins located at the distal tip of the tail fibers of diverse T-even phages and has been shown to determine the host specificity in phages ^71^. The adhesins encoded by hafunaviruses form four distinct phylogenetic groups (Groups 1 to 4), with the corresponding viruses displaying distinct infectivity patterns on the tested haloarchaeal strains (Fig. 5, Fig. S7). This is most prominent in the case of viruses encoding the largest adhesin groups, Group 1 and Group 3. For instance, four strains, *Halorubrum* sp. SS8-7, *Halorubrum sodomense, Haloarcula californiae* and *Halobellus* sp. SS6-7, are particularly sensitive to viruses encoding Group 1 adhesins, with the latter two strains being exclusively infected by viruses encoding this group of adhesins. Conversely, no virus of this group is able to infect *Halorubrum* strains SS7-4, SS10-3 and SS5-4 or *Haloarcula* sp. SS8-5. Nearly all viruses encoding Group 3 adhesins are able to infect *Halorubrum* sp. SS9-12, whereas *Halorubrum* sp. SS6-1 and *Haloterrigena* sp. S13-7 are completely insensitive to this group of viruses (Fig. 5) (see Supplementary text for details). In all *Hafunaviridae* species, the adhesin gene is located within a hypervariable region (Fig. S6), which has been previously suggested to encode tail fiber proteins ^22^. Collectively, our results implicate the adhesin-encoding gene as the key host range determinant in hafunaviruses. Importantly, while the host range pattern can be explained, even if partly, by the adhesin phylogeny, overall intergenomic relationships between hapunaviruses is a poor predictor of the host range. Indeed, viruses belonging to the same species (i.e., >95% genome-wide identity) encode adhesins from different phylogenetic groups and display distinct host ranges.

**Fig. 5.**
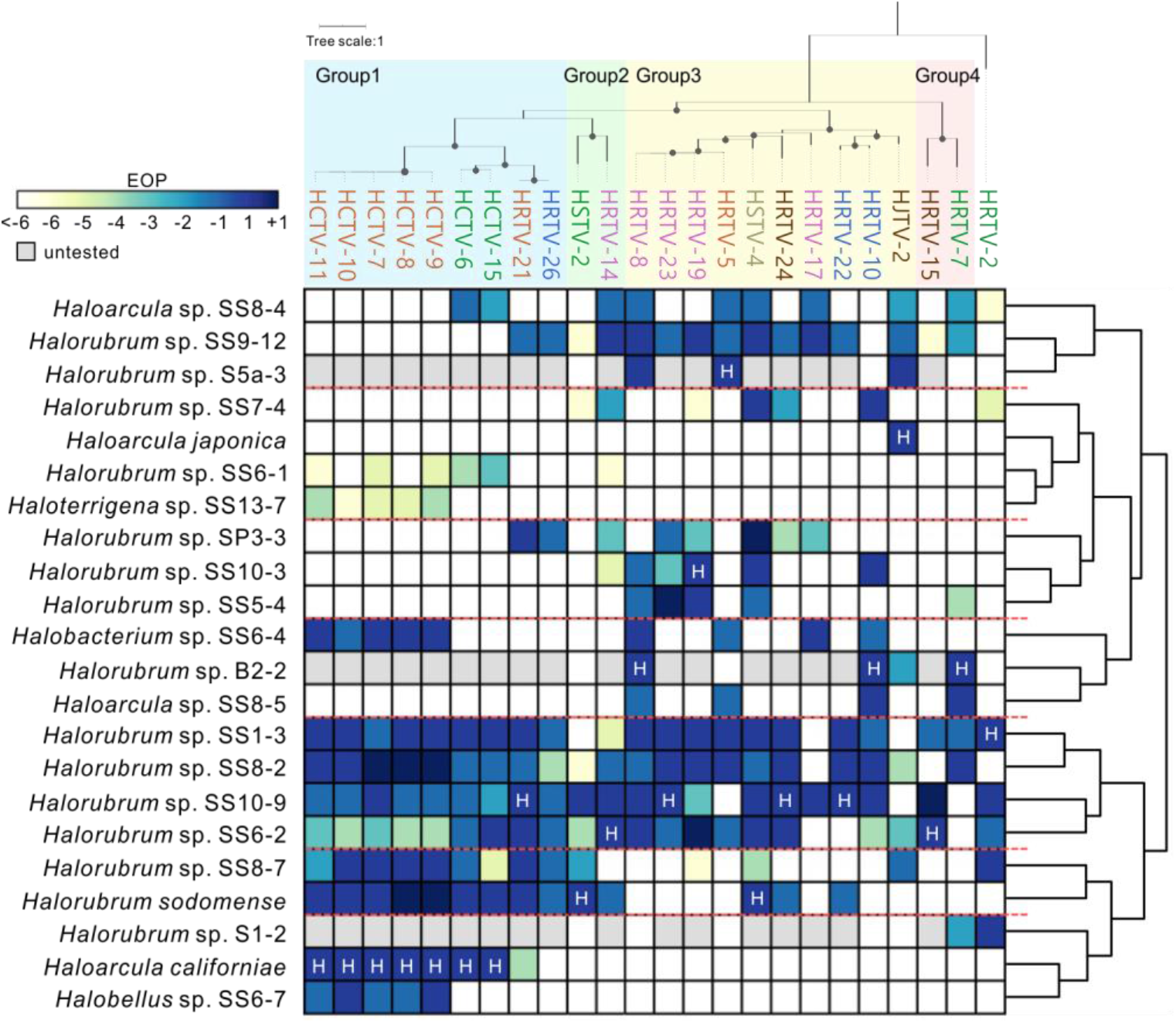
Host ranges of hafunaviruses. The heatmap shows the efficiency of plating (EOP) of hafunavirus isolates on different haloarchaeal strains. The EOP of the virus on its original isolation host has been set to one and marked with H. Number 1 is equal to the EOP on the original host. Numbers −1, −2,… refer to 10^−1^, 10^−2^,… and +1 refers to 10^1^ when compared to the EOP on the original host, as shown in Table S9. The isolation host for HRTV-26 is *Halorubrum* sp. SS13-13, which occasionally did not form a lawn and was not included in this analysis. The titer of HRTV-26 on *Halorubrum* sp. SS13-13 was 2 × 10^8^ pfu/ml, which was considered here as one and used for comparison. The hosts were clustered based on the similarity of EOPs of the tested viruses, as represented by the dendrogram on the right side. The upper panel shows the maximum likelihood phylogenetic tree of the tail fiber adhesin proteins encoded by the analyzed viruses. The adhesin of HRTV-2, the amino acid sequence of which is markedly divergent from those of other hafunaviruses, is set as the outgroup. The four distinct groups of adhesins are displayed with colored blocks. Bootstrap values greater than 90% are indicated in the nodes by dots. Virus names are colored according to the species to which they belong.

### Continuity between archaeal tailed viruses across biomes

The previous study of viral communities using metagenomics approach revealed abundant presence of tailed viruses in hypersaline environments ^72^. To assess the extent of sequence diversity and environmental distribution of halophilic arTVs and to explore the evolutionary relationships between tailed viruses from different environments, we searched the Integrated Microbial Genomes and Microbiomes (IMG/M) database for homologs of MCP, portal and the large terminase subunit (TerL) proteins representing each family of arTVs. The searches collectively yielded 146,679 contigs encoding at least one of the three viral hallmark proteins. The dataset was supplemented with reference sequences from viruses for which hosts are known or predicted *in silico*, including tailed (pro)viruses associated with methanogens, MGI and MGII Euryarchaeota, Thaumarchaeota, Thermoplasmata and Aigarchaeota ^26,27,29,31-35^. In addition, sequences of uncultivated haloviruses previously obtained through fosmid sequencing from the Santa Pola saltern, Spain, were also added to the dataset ^73^. Among the three hallmark proteins, TerL appeared to be the least specific to archaeal viruses, likely due to its relatively high conservation across all tailed bacterial and archaeal viruses. Accordingly, searches queried with haloarchaeal virus TerL against the IMG/M database retrieved a high number of significant hits from diverse environments (Fig. S10). By contrast, MCP and portal proteins were more specific to halophilic arTVs and retrieved a higher ratio of homologs from hypersaline ecosystems compared to those from other habitats. This is consistent with the fact that haloviral MCPs and portal proteins typically display low sequence similarity to their bacteriophage homologs in BLASTP searches. Consequently, MCP and portal proteins were used as specific markers for arTVs.

The MCP tree splits into nine well-supported clades, each represented by haloarchaeal tailed virus isolates from one or more families (Fig. 6). A similar result was obtained with the portal protein homologs (Fig. S10). Almost all clades in the MCP tree contain numerous homologs originating from hypersaline environments (Fig. 6), with the corresponding metagenomes derived from all seven continents, testifying to the wide geographic distribution of the halophilic arTVs. This result suggests that the haloarchaeal virus isolates described herein adequately represent the overall diversity of haloarchaeal tailed virus communities.

**Fig 6.**
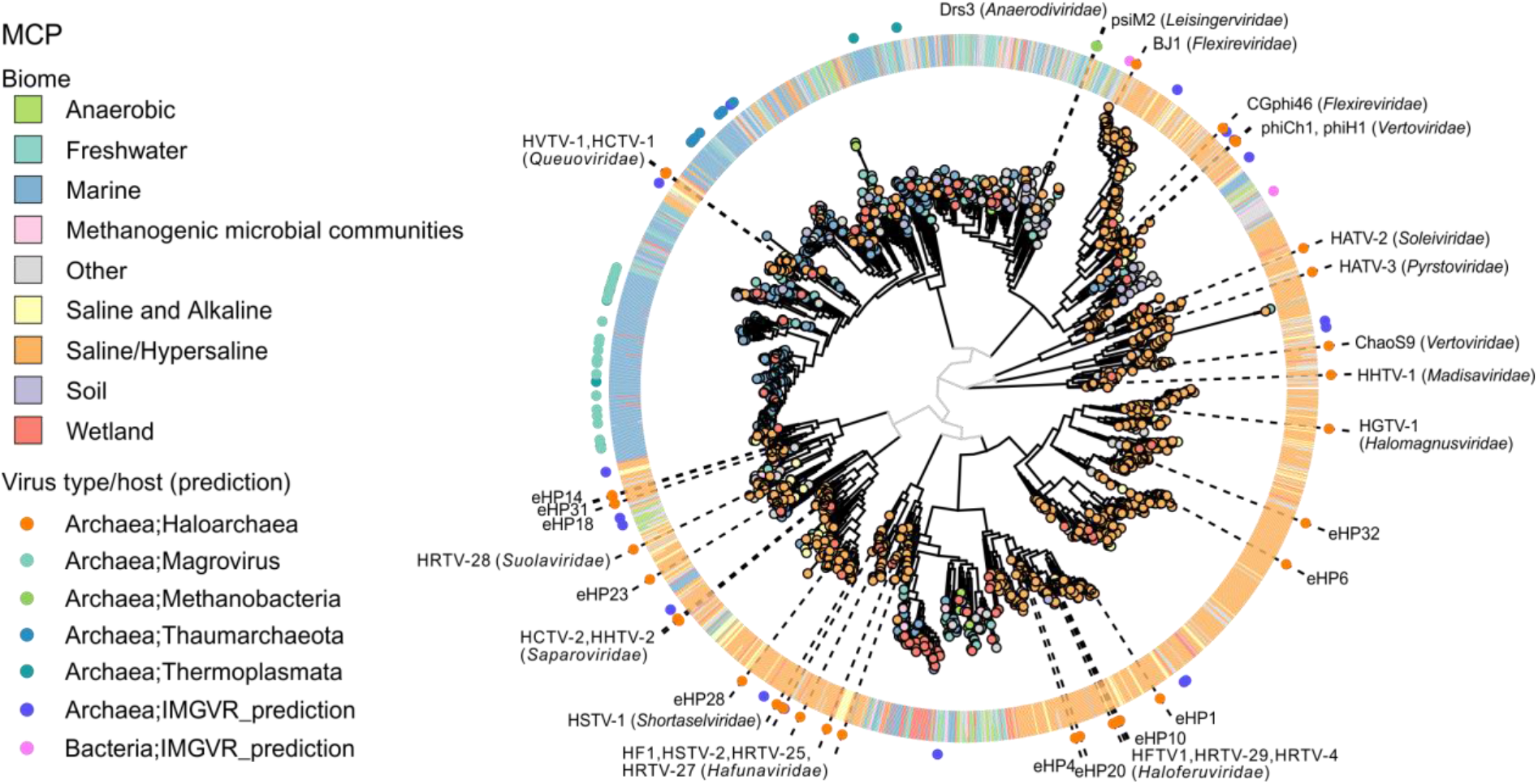
Phylogenetic reconstruction of the MCPs encoded by arTVs. The homologs retrieved from metagenomes were clustered at 0.9 protein sequence identity and the representative of each cluster is indicated as a circle at the tip of the branch with the color reflecting the corresponding biome (see the legend on the left). The ring outside of the tree consists of tiles indicating the biome compositions from the corresponding protein cluster. The dot in the outermost ring indicates the (predicted) virus type in the corresponding branch and is colored based on the host predicted. The names of the arTV isolates of haloarchaea and methanogens as well as the fosmids generated from a saltern of Santa Pola in Spain ^73^ are shown on the tree. The virus family names are indicated next to the corresponding arTV isolates.

Interestingly, although searches with the hallmark proteins were performed against the global IMG/M database, only a handful of sequences affiliated to bacteria/phages were retrieved (Fig. 6, S10). By contrast, haloarchaeal viruses formed well-supported clades with the previously characterized viruses associated with marine and methanogenic archaea, suggesting that all arTVs are more closely related to each other than to tailed bacteriophages, consistent with the GRAViTy analysis (Fig. 1). Indeed, three of the clades in the MCP tree contained considerable fractions of taxa originating from non-hypersaline environments (Fig. 6). The largest of these clades is dominated by MCP homologs from marine environments, with a relatively small subclade including haloarchaeal viruses of the *Queuoviridae* (e.g. HVTV-1), interspersed between reference viruses associated with MGII Euryarchaeota (magroviruses) ^34,35^, marine Thaumarchaeota ^32^ and Thermoplasmata ^33^ (Fig. 6). A similar clustering is also retrieved in the portal protein tree (Fig. S10), suggesting that the ancestor of the *Queuoviridae* has evolved from the midst of marine archaeal viruses.

The second MCP tree clade containing sequences from diverse environments is represented by viruses from the *Flexireviridae* and *Vertoviridae* (e.g. BJ1 and phiCh1, respectively; note that the MCP of ChaoS9 from the *Vertoviridae* clusters with that of HHTV-1 from the *Madisaviridae*). This group of halovirus MCPs is embedded within an assemblage of homologs from diverse non-saline environments, including anoxic habitats (Fig. 6). Consistently, the clade includes viruses psiM2 and Drs3 (families *Leisingerviridae* and *Anaerodiviridae*, respectively) infecting methanogenic archaea (Fig. 6). The same relationship is recapitulated in the portal protein tree (Fig. S10). Therefore, viruses from the families *Flexireviridae* and *Vertoviridae* appear to have evolved from viruses inhabiting moderate, non-saline environments, although the exact evolutionary history remains obscure, given the complicated composition and relatively poor sampling of this clade.

Finally, the third MCP tree clade is represented by viruses from the families *Haloferuviridae* and *Halomagnusviridae* (e.g. HFTV1 and HGTV-1, respectively). Unlike in the other two cases, where haloarchaeal viruses were nested among taxa from non-hypersaline environments, in this clade, we observe a reverse situation. Namely, a group of sequences, largely originating from anoxic environments, such as methanogenic microbial communities, anaerobic digesters, wetlands, etc., with at least one contig predicted to represent a virus infecting a methanogenic archaeon, forms a sister clade to haloviruses of the *Haloferuviridae*, and is further embedded within a clade dominated by sequences from hypersaline environments (Fig. 6). A parsimonious explanation for such clustering involves evolution of a group of methanogenic archaeal viruses from within the diversity of haloarchaeal viruses.

Collectively, these results suggest that there is a continuity between arTVs across different biomes. At the same time, however, it becomes clear that haloarchaeal viruses represent a polyphyletic assemblage, with several groups immigrating into hypersaline environments from other habitats with lower salinity. Presumably, the ancestors of these groups gained the ability to infect halophilic archaea by accumulating mutations within the tail fiber genes, akin to those described above for hafunaviruses. We note that natural mixing between marine and hypersaline ecosystems (e.g., through evaporation of coastal marine water) provides an ecological setting for marine viruses to encounter halophilic archaea. Thus, hypersaline environments, once considered as an isolated extreme environment ^74,75^, emerge as an integral part of the whole ecosystem.

## CONCLUDING REMARKS

In the present study, we sequenced 37 haloarchaeal tailed virus isolates which originate from four spatially remote locations, more than doubling the current number of sequenced arTVs (Table S1). Through a systematic analysis of all 63 available arTVs, we propose 14 new families for their classification. This sets the first framework for the taxonomy of arTVs and it is justified to associate them with the class of *Caudoviricetes* with the tailed bacteriophages due to their shared evolutionary roots. Half of the arTV families are currently represented by a single virus isolate, indicating the scarce sampling of haloarchaeal virome.

It has been previously shown that arTVs share virion architecture, assembly principles and genome organization with their bacterial virus counterparts ^11,13,30,38,76^. In this study, we focused on metabolic genes and showed that arTVs carry a rich repertoire of counterdefense and metabolic genes, predominantly those involved in DNA and RNA metabolism. Many of these genes are also commonly encoded by tailed bacteriophages, suggesting common propagation strategies for these bacterial and archaeal viruses. Nevertheless, some of the identified metabolic genes have never been reported in other viruses, e.g. *carA* which is predicted to promote the de novo synthesis of pyrimidines, RNA methyltransferase activator *trm112* implicated in modulation of protein translation, and the gene encoding a putative ribosomal protein L21e.

Myoviruses of the *Hafunaviridae*, the most populous family of arTVs, have broad host ranges, with strains of the same species displaying considerable variation in the host range. We found that the host range specificity in these viruses is primarily determined by mutations within a gene encoding a tail adhesin protein, which resembles that of T-even bacteriophages ^71,77^. High sequence divergence of adhesins encoded by hafunaviruses likely allows these viruses to explore a broad landscape of receptor molecules on the host surface, boosting the competitiveness of this group of viruses in the environments with some of the highest reported virus-to-host ratios. Despite the fundamental differences of archaeal and bacterial cell surface structures ^78^, the host range determinant of tailed viruses, the tail adhesins, can be highly similar in sequence features between arTVs and tailed bacteriophages. This highlights the versatility of the tail fiber structure towards diverse host receptors, which may underlie the success of the *Caudoviricetes* viruses.

Finally, the survey of metagenomic databases using haloviral hallmark proteins uncovered a considerable diversity and global distribution of arTVs. Remarkably, our results point to the polyphyletic origins of the arTVs, with some of the groups originating from archaeal viruses associated with marine and methanogenic hosts, indicative of virus movement across different biomes. Given that in hypersaline environments, the arriving viruses are exposed to extremely high salt concentrations both inside and outside of the cell ^79^, the adaptation must have entailed a substantial evolution of the virion proteins. Indeed, some of the halophilic arTVs have been shown to be non-infectious under low salt conditions ^11,37^. Remarkably, however, the effect was reversible and upon addition of salt, the infectivity was restored, suggesting that virions of haloarchaeal viruses can maintain stability and integrity under a wide range of environmental conditions. Further sampling and isolation of arTVs, especially those infecting non-halophilic archaea, will be important for further understanding the ecology and evolution of this important group of viruses.

## MATERIALS AND METHODS

### Preparation of virus samples

Viruses grown in this study had been isolated previously ^16,17^. The host strains used for virus production are listed in Table S1. Viruses and strains were grown aerobically at 37°C in modified growth medium (MGM) made using artificial salt water (SW) (23% (w/v) in broth, 20% in solid, and 18% in top agar) (http://www.haloarchaea.com/resources/halohandbook/). The double-layer plaque assay method was used for virus growth and quantification, and virus stocks were prepared from semi-confluent plates as previously described ^16,17^. Viruses were precipitated from stocks with polyethylene glycol 6000 (final concentration of 10% (w/v)) by magnetic stirring for 1 h at 4°C, collected by centrifugation (Fiberlite F14 rotor, 9820 g, 40 min, 4°C), and resuspended in 18% SW. Virus samples were purified by rate zonal centrifugation in 5-20 % (w/v) sucrose (18 % SW; Sorvall AH629 rotor, 103586 g, 40 min, 15°C), concentrated by differential centrifugation (Sorvall T647.5 rotor, 113580 g, 3 h, 15°C) and resuspended in 18% SW.

### Viral genome isolation, sequencing and annotation

Nucleic acids were extracted from purified virus samples either using the PureLink Viral RNA/DNA Mini Kit (Thermo Fisher Scientific) or by phenol/ether method combined with ethanol precipitation. The viral genome libraries were prepared with the Nextflex PCRFree kit (Bioo Scientific), and sequenced by Illumina Miseq (Illumina, San Diego, CA) with paired-end 250-bp read length (Biomics Platform, Institut Pasteur, France). Sequenced reads were quality-trimmed using fqCleaner v0.5.0 and assembled using clc_assembler v4.4.2 implemented in Galaxy-Institut Pasteur. The genomic termini and packaging mechanism were determined by PhageTerm ^80^. ORFs were predicted using RAST v2.0 server ^81^, and tRNA genes using tRNAscan-SE ^82^. Functional annotations of putative genes were performed using HHpred ^51^. The average nucleotide identity between viruses was calculated by ANI Calculator ^83^, whereas the global alignment of amino acid sequences was carried out by EMBOSS Needle tool ^84^. Multiple sequence alignments were performed by Promals3D ^85^.

### Taxonomy assignment

The viral genomes were subjected to GRAViTy ^47,48^ (http://gravity.cvr.gla.ac.uk) and vConTACT 2.0 ^49^ for taxonomy assignment. We incorporated the annotated genomes of arTVs into the genomic datasets which were (i) built for bacterial and archaeal dsDNA viruses from the ICTV 2016 Master Species List 31V1.1 for GRAViTy analysis, and (ii) embedded in vConTACT 2.0 with the NCBI Bacterial and Archaeal Viral RefSeq V88 and default settings for analysis. The virus network generated by vConTACT 2.0 was visualized with Cytoscape software v.3.7.2 (https://cytoscape.org/), using an edge-weighted spring-embedded model that genomes sharing more PCs were positioned closer to each other. Pair-wise genomic comparison was performed by EasyFig with the E-value cutoff of 1 × 10^−3^ and minimum protein length of 15 residues ^86^. The overall similarity among arTVs on genus and family levels were analyzed and proteins with over 30% amino acid sequence identity and E-value < 1 × 10^−25^ in our arTV database were counted as homologous. The results were visualized with Circos plot ^87^, whereas Box plots were prepared using OriginLab v9.8 (OriginLab. Inc., USA).

### Virus-host interactions

Virus-host pairs tested for hafunaviruses in this study are shown in Table S9. For initial screening of virus-host interactions, 10-µl drops of undiluted and 100-fold diluted virus stocks were spotted on the lawn of early-stationary growing strains prepared by the double-layer method, and 10-µl drops of MGM broth were spotted as negative control. After 2–5 days of incubation, plates were screened for the growth inhibition. When inhibition was observed, plaque assay was used to verify virus-host interactions and quantify the EOP. The titers above 10^3^ PFU/mL could be detected. Data were normalized by comparing the EOP to that on the original host strain (set as 1). For phylogenetic analyses of hafunavirus adhesins, protein sequences were aligned with Promals3D ^85^ and the maximum likelihood phylogenetic tree was constructed using PhyML 3.0 ^88^ with the best-fitting model identified by the program being WAG +G +F. The heatmap of the virus host ranges was generated based on the EOP results and clustered according to the phylogeny of viral adhesins as well as EOP similarity of the tested hosts using pheatmap package in R.

### Metagenomic database screening

The MCP, portal and TerL protein sequences of the isolated halophilic arTVs were used as queries in a BLAST search of 31,344 public metagenomes available in the Integrated Microbial Genomes and Microbiomes (IMG/M) database at JGI with a threshold of 100 bit score and 80% sequence coverage. Sequences of the three markers from available (pro)viruses of methanogen, marine group I and group II Euryarchaeota, Thaumarchaeota, Thermoplasmata, Aigarchaeota and fosmids from a saltern of Santa Pola in Spain were extracted ^26,27,29,31-35,72^. These sequences were subjected to BLAST searches against our retrieved metagenomic sequences with the same threshold as above and were used as reference sequences in the phylogenetic analysis. Proteins from different collection origins were clustered at 90% sequence identity separately and joint as the final datasets. Sequences in the datasets of the three hallmark proteins were aligned using MAFFT program v7 ^89^, followed by removal of poorly aligned positions by trimAl with gap threshold of 0.2 ^90^. The phylogenetic trees were constructed using a JTT+CAT model by FastTree (version 2.1.11) ^91^, and visualized using ggtree v2.4.1 ^92^. Biome information for IMG/M metagenomes were obtained from the Gold database ^93^. All IMG/M contigs with a hit to an arTV marker were also cross-referenced with contigs in the IMG/VR database ^94^. For contigs detected as viral and included in IMG/VR, the host prediction was obtained when available at the domain level (i.e. bacteria vs archaea) and displayed on the trees.

## ACKNOWLEDGEMENTS

We thank Sari Korhonen, Matti Ylänne, and Carlos Lapedriza for skillful technical assistance. This work was supported by l’Agence Nationale de la Recherche grant ANR-20-CE20-0009-02 (to M.K.) and the European Union’s Horizon 2020 research and innovation program under grant agreement 685778, project VIRUS X (to D.P.). Y.L. is a recipient of the Pasteur-Roux-Cantarini Fellowship from Institut Pasteur. The Ella and Georg Ehrnrooth Foundation and the Finnish Cultural Foundation are sincerely acknowledged (grants to T.D.). The facilities and expertise of the HiLIFE Biocomplex unit at the University of Helsinki, a member of Instruct-ERIC Centre Finland, FINStruct, and Biocenter Finland are gratefully acknowledged. The work conducted by the U.S. Department of Energy Joint Genome Institute (S.R.) is supported by the Office of Science of the U.S. Department of Energy under contract no. DE-AC02-05CH11231. The development and application of GRAViTy analysis was supported by a grant to PS from the Wellcome Trust (WT108418AIA).

## Notes

### Competing Interest Statement

The authors have declared no competing interest.

